# Diversified repertoire of phage resistance in *Klebsiella pneumoniae* and bidirectional steering effects impacting antibiotic susceptibility

**DOI:** 10.1101/2022.11.14.516531

**Authors:** Sue C. Nang, Jing Lu, Heidi H. Yu, Hasini Wickremasinghe, Mohammad A. K. Azad, Meiling Han, Jinxin Zhao, Gauri Rao, Phillip J. Bergen, Tony Velkov, Norelle Sherry, David T. McCarthy, Saima Aslam, Robert T. Schooley, Benjamin P. Howden, Jeremy J. Barr, Yan Zhu, Jian Li

**Affiliations:** Department of Microbiology, Biomedicine Discovery Institute, Monash University, Clayton, Victoria, Australia; Division of Pharmacotherapy and Experimental Therapeutics, Eshelman School of Pharmacy, University of North Carolina, Chapel Hill, North Carolina, USA; Biomedicine Discovery Institute, Department of Pharmacology, Monash University, Clayton, Victoria, Australia; Microbiological Diagnostic Unit Public Health Laboratory, Department of Microbiology and Immunology, The University of Melbourne at The Peter Doherty Institute for Infection and Immunity, Melbourne, Victoria, Australia; Department of Civil Engineering, Monash University, Clayton, Victoria, Australia; Division of Infectious Diseases and Global Public Health, Department of Medicine, University of California San Diego, La Jolla, California, USA; School of Biological Sciences, Monash University, Clayton, Victoria, Australia; Tianjin Institute of Industrial Biotechnology, Chinese Academy of Sciences, Tianjin, China

## Abstract

Bacteriophage (phage) therapy is rising as a promising anti-infective option to combat antimicrobial resistance; however, its clinical utilization is severely hindered by the potential emergence of phage resistance. Fortunately, certain phage resistance mechanisms can restore bacterial antibiotic susceptibility, making the combination of phages with antibiotics a potential strategic approach. Here, we demonstrated that phage resistance can also lead to increased antibiotic resistance and provided mechanistic insights into bacterial phage defense mechanisms. We discovered a repertoire of phage resistance mechanisms in *Klebsiella pneumoniae*, including the disruption of phage binding site (*fhuA*::Tn and *tonB*::Tn), extension of phage latent period (*mnmE*::Tn and *rpoN*::Tn) and increased mutation frequency (*mutS*::Tn and *mutL*::Tn). Different from the prevailing view that phage resistance re-sensitizes antibiotic-resistant bacteria, we revealed a bidirectional steering effect on the bacterial antibiotic susceptibility. Specifically, it was uncovered that, while *rpoN*::Tn became more susceptible to colistin, *mutS*::Tn and *mutL*::Tn caused increased resistance to rifampicin and colistin. Our findings highlight the diversified strategies utilized by *K. pneumoniae* to overcome phage infection and the parallel effect on the antibiotic susceptibility. Mechanism-guided phage steering represents a rational strategy that should be incorporated into phage therapy to better inform clinical decisions.

## Introduction

Bacteriophages (phages) are viruses that specifically target bacteria (1, 2). The earliest recorded therapeutic practice of phages for the treatment of bacterial infections took place almost a century ago, prior to the discovery of penicillin (3–6). As the golden age of antibiotic was on the rise, phage therapy was abandoned, mainly due to technical and logistical hurdles (3). In recent times, the ‘perfect storm’ of widespread antimicrobial resistance (AMR) combined with the sparse antibiotic development pipeline has dramatically reduced the antibiotic armamentarium available to physicians for treating multidrug-resistant (MDR) infections (7, 8). Of particular concern are the ESKAPE pathogens which include *Klebsiella pneumoniae*, that possess a great ability to ‘escape’ the antibacterial effects of many antibiotics through a wide range of resistance mechanisms (9, 10). With the dawn of a post-antibiotic era rapidly emerging, the World Health Organization (WHO) and the Centers for Disease Control and Prevention (CDC) have declared AMR as a major global health threat (11, 12). Clearly, alternative antimicrobial therapies are urgently required and therefore, significant interest has risen in the use of phages for treating MDR bacterial infections (3, 4, 13). To date, the clinical utility of phages has been hampered by a poor knowledge base on their biology, resistance mechanisms and combination use with antibiotics (3).

Phage steering has recently received significant favorable attention in the AMR field, involving the use of phages to kill susceptible bacterial cells while re-sensitizing the remaining phage-resistant bacterial population to antibiotics (14, 15). The utility of phage steering lies in the prospect of eliminating the entire bacterial population using combinations of phages and antibiotics; however, the intricacies and limitations of phage steering have yet to be explored.

Improving the knowledge base to guide phage steering based on underlying mechanisms is of paramount importance to inform clinical decisions on phage and antibiotic selections. Here, we discovered an unprecedented bidirectional phage steering effect with both decreased and increased antibiotic susceptibility associated with different phage resistance mechanisms in *K. pneumoniae*. Our findings may have significant implications in both basic and clinical biomedical science on phage therapy, ultimately advancing its application in patient care.

## Results

### Phage resistome screening reveals involvement of diverse genes

Utilizing a strictly lytic *Siphoviridae* phage pKMKP103_1 (**Figure 1**), we conducted a genome-wide resistome study by investigating the susceptibility of individual transposon mutants of *K. pneumoniae* MKP103 (sequence type 258, capsule type K107) (16). Screening of phage resistome identified six genes, *fhuA*, *tonB*, *mnmE*, *rpoN*, *mutS* and *mutL*, in *K. pneumoniae* MKP103 that are required for effective pKMKP103_1 infection (**Figure 2a and Table S1**). Subsequently, we proceeded to confirm the phage susceptibility profile of these candidate transposon mutants by evaluating the ability of pKMKP103_1 to form clear plaques on soft agar containing the respective mutants (**Figure 2b**), and the killing kinetics of pKMKP103_1 against the respective mutants in broth (**Figure 2c**). Against *fhuA*::Tn and *tonB*::Tn, pKMKP103_1 did not show any lysis zone on agar plates (**Figure 2b**) nor killing in broth (**Figure 2c**), potentially attributed by the inability of pKMKP103_1 to infect these mutants. The efficiency of plating (EOP) of pKMKP103_1 on *mnmE*::Tn was reduced by 10,000-fold in comparison to the wild-type (WT) (**Figure 2b**), showing that the ability of pKMKP103_1 to form visible plaque on this mutant was restricted. A reduced bacterial killing was also illustrated by *mnmE*::Tn with at least 1,000-fold lower bacterial reduction at 1 h (**Figure 2c**). Interestingly, although the EOP of pKMKP103_1 on *rpoN*::Tn was unaffected (**Figure 2b**), no bacterial killing was observed over 24 h when cultured in broth (**Figure 2c**). With *mutS*::Tn and *mutL*::Tn, the EOP of pKMKP103_1 was similar to the WT (**Figure 2b**); however, a weaker initial killing activity was evident in broth (**Figure 2c**). The restoration of the phenotype of these mutants to that of the WT following complementation confirmed the involvement of these individual genes for effective pKMKP103_1 infection.

**Figure 1:**
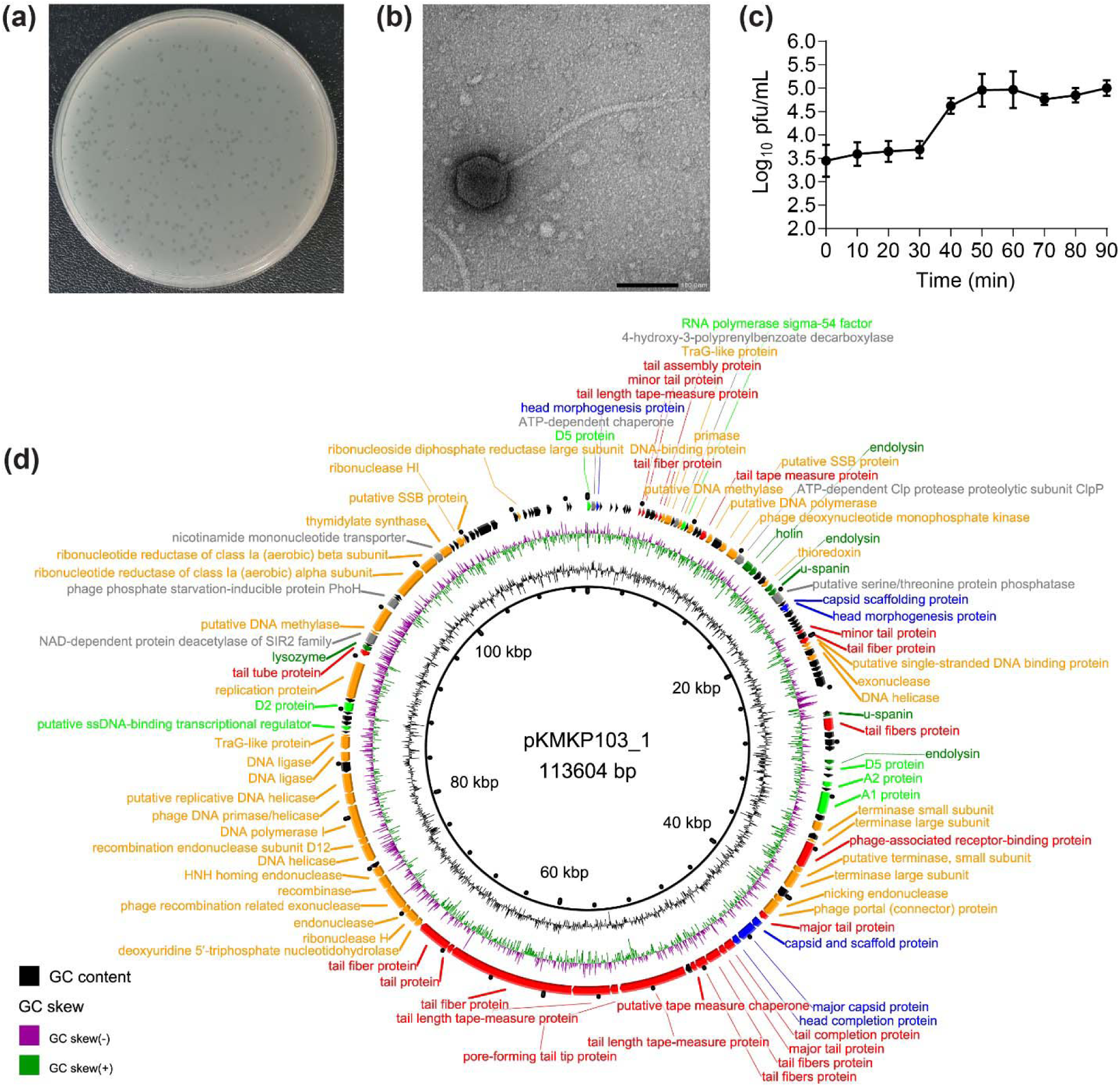
Characterization of phage pKMKP103_1. **(a)** Plaque morphology of pKMKP103_1 plated on *K. pneumoniae* MKP103. **(b)** A representative TEM image of pKMKP103_1. The scale bar represents 100 nm. **(c)** One-step growth curve of pKMKP103_1 with *K. pneumoniae* MKP103. Data are presented as mean ± SD (*n* = 3). **(d)** Visualization of pKMKP103_1 genome. The colors represent the functions of the genes: orange (DNA packaging, replication, and modification), lime (transcription), green (host lysis), blue (head structural components), red (tail structural components), gray (others/unknown functions), and black (hypothetical).

**Figure 2:**
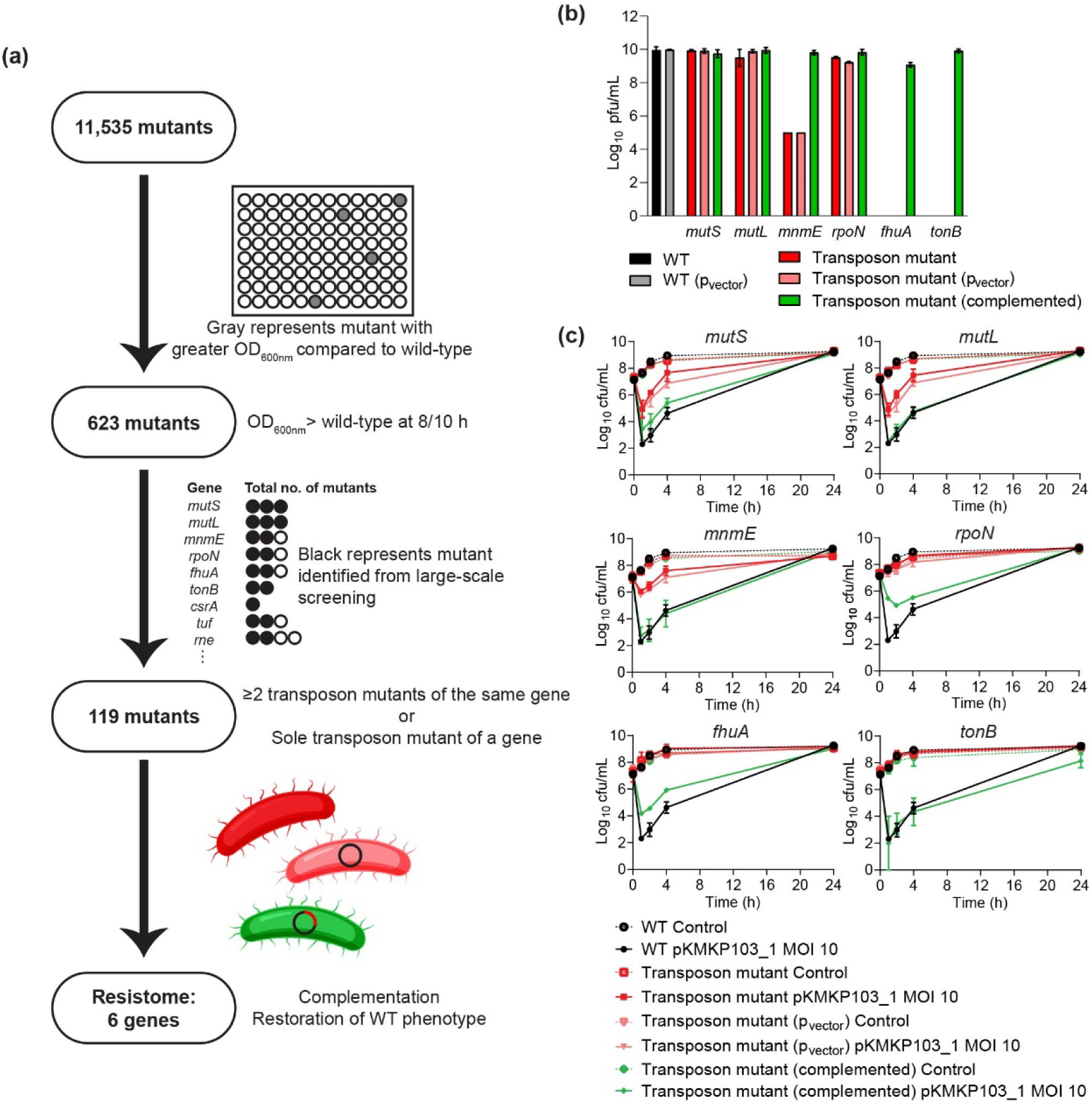
Identification of *K. pneumoniae* MKP103 genes that are correlated with phage pKMKP103_1 resistance and the associated resistance phenotype. **(a)** Workflow for the genome-wide transposon screening of pKMKP103_1 resistome using the transposon mutant library. **(b)** The plaque forming unit (pfu) when pKMKP103_1 was plated on the respective *K. pneumoniae* MKP103 strains. Individual plaque counts were not available for *mnmE*::Tn and *mnmE*::Tn (p_vector_); the count was calculated based on the highest dilution in which lysis zone could be observed. Data are presented as mean ± SD (*n* = 3). **(c)** Time-kill kinetics of pKMKP103_1 against *K. pneumoniae* MKP103 and mutant strains at an MOI of 10. Data are presented as mean ± SD (*n* = 3).

### Phage resistance occurred with multiple mechanisms

With different phage susceptibility profiles observed for the aforementioned six mutants, we proceeded to investigate the mechanisms causing these differential degrees of resistance. The adsorption of pKMKP103_1 to the bacterial surface of *fhuA*::Tn and *tonB*::Tn was completely inhibited (**Figure 3a–b**). As the adsorption of pKMKP103_1 onto the bacterial surface of *mnmE*::Tn, *rpoN*::Tn, *mutS*::Tn and *mutL*::Tn was unaffected (**Figure 3b**), we subsequently examined the impact for the disruption of these genes on the lytic lifecycle of pKMKP103_1 by examining the one-step growth profiles of the phage with respective mutants. With either *mutS*::Tn or *mutL*::Tn as the host strain, similar one-step growth profiles of pKMKP103_1 to that observed with the WT were demonstrated (**Figure 3c**). Interestingly, a longer latent period (40 min) compared to the WT (30 min) was displayed by pKMKP103_1 when *mnmE*::Tn and *rpoN*::Tn were employed as the host strain; however, there were no substantial differences in the burst size (**Figure 3c**). Given that bacterial isolates with interrupted *mutS* and *mutL* are commonly known as hypermutators (17–20), we investigated the impact of the disruption of these genes on the bacterial mutation rate towards pKMKP103_1. Compared to the WT (7.4 ± 4.7 × 10^-6^), a significant increase in mutation frequency to pKMKP103_1 was discovered with *mutS*::Tn (1.4 ± 1.1 × 10^-3^) and *mutL*::Tn (1.6 ± 1.7 × 10^-3^) (**Figure 3d**). Complementation of *mutL* completely restored the mutation frequency of *mutL*::Tn (7.3 ± 3.3 × 10^-6^). While for *mutS*::Tn, a reduced mutation frequency (1.8 ± 2.1 × 10^-4^) was observed following complementation; however, it did not reach the WT level.

**Figure 3:**
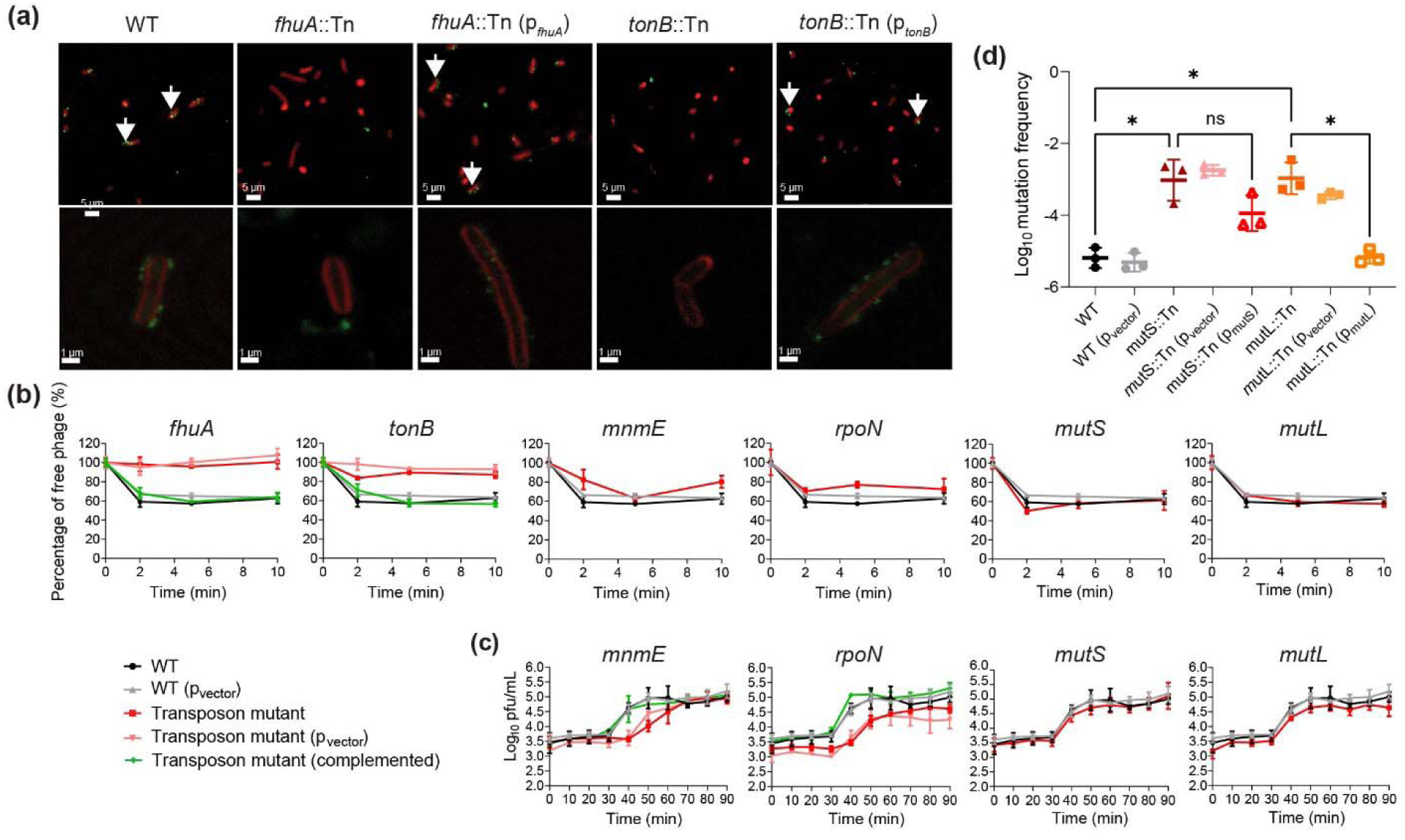
Phage pKMKP103_1 adsorption, one-step growth profiles and bacterial mutation frequency towards pKMKP103_1. **(a)** Visualization of pKMKP103_1 adsorption determined using fluorescent (top panels) and confocal (bottom panels) microscopy. The pKMKP103_1 was stained with SYBR^TM^ Gold strain (green) and *K. pneumoniae* MKP103 strains were stained with FM^TM^ 4-64FX (red). **(b)** pKMKP103_1 adsorption profile with respective *K. pneumoniae* strains (*n* = 3). **(c)** One-step growth profile of pKMKP103_1 in the presence of different *K. pneumoniae* strains. Data are presented as mean ± SD (*n* = 3). **(d)** The mutation frequency of *K. pneumoniae* strains to phage pKMKP103_1. Data are presented as mean ± SD (*n* = 3). Statistical significance is denoted with \**p* <0.05.

### Bidirectional steering effects on antibiotic susceptibility in phage-resistant mutants

Phage steering has been widely portraited as a promising antibacterial strategy, having the potential to eliminate the entire bacterial population with phage-antibiotic combinations (14, 15, 21). Hereby, we examined the impact of different resistance mechanisms identified earlier on the antibiotic susceptibility of *K. pneumoniae* MKP103 (**Figure 4a and Table S2**). In comparison to the WT (rifampicin minimum inhibitory concentration [MIC]=64 mg/L; colistin MIC=32 mg/L), phage-resistant *K. pneumoniae* mutants with the hypermutator phenotype (*mutS*::Tn and *mutL*::Tn) exhibited increased resistance to rifampicin (MIC>128 mg/L) and colistin (MIC=128 mg/L); whereas *rpoN*::Tn showed decreased resistance to colistin (MIC=4 mg/L; **Figure 4a**). The complementation of mutants with respective WT genes restored the antibiotic susceptibility to the WT level, confirming that the disruption of these genes was indeed directly associated with the change in bacterial susceptibility to rifampicin and/or colistin in *K. pneumoniae* MKP103 (**Figure 4a**). Moreover, we conducted population analysis profiles (PAPs) of colistin with *mutS*::Tn, *mutL*::Tn and *rpoN*::Tn to further quantify the susceptibility change. The PAPs revealed an approximately 100-fold increase in the subpopulation of *mutS*::Tn and *mutL*::Tn on Mueller Hinton (MH) agar containing 128 mg/L colistin, when compared to the WT (**Figure 4b**). In the case of *rpoN*::Tn, compared to the WT, there was an approximately 1,000-fold reduction in the subpopulation growing on MH agar in the presence of 16 and 32 mg/L colistin and approximately 100-fold reduction in the subpopulation growing with 64 mg/L colistin (**Figure 4b**). To examine the association between lipid A modifications and colistin susceptibility (22), our lipid A profiling revealed that, compared to the WT, there were decreased abundance of 4-amino-4-deoxy-L-arabinose (L-Ara4N)-modified lipid A and increased abundance of 2-hydroxymyristate lipid A in *rpoN*::Tn (**Figure 4c**).

**Figure 4:**
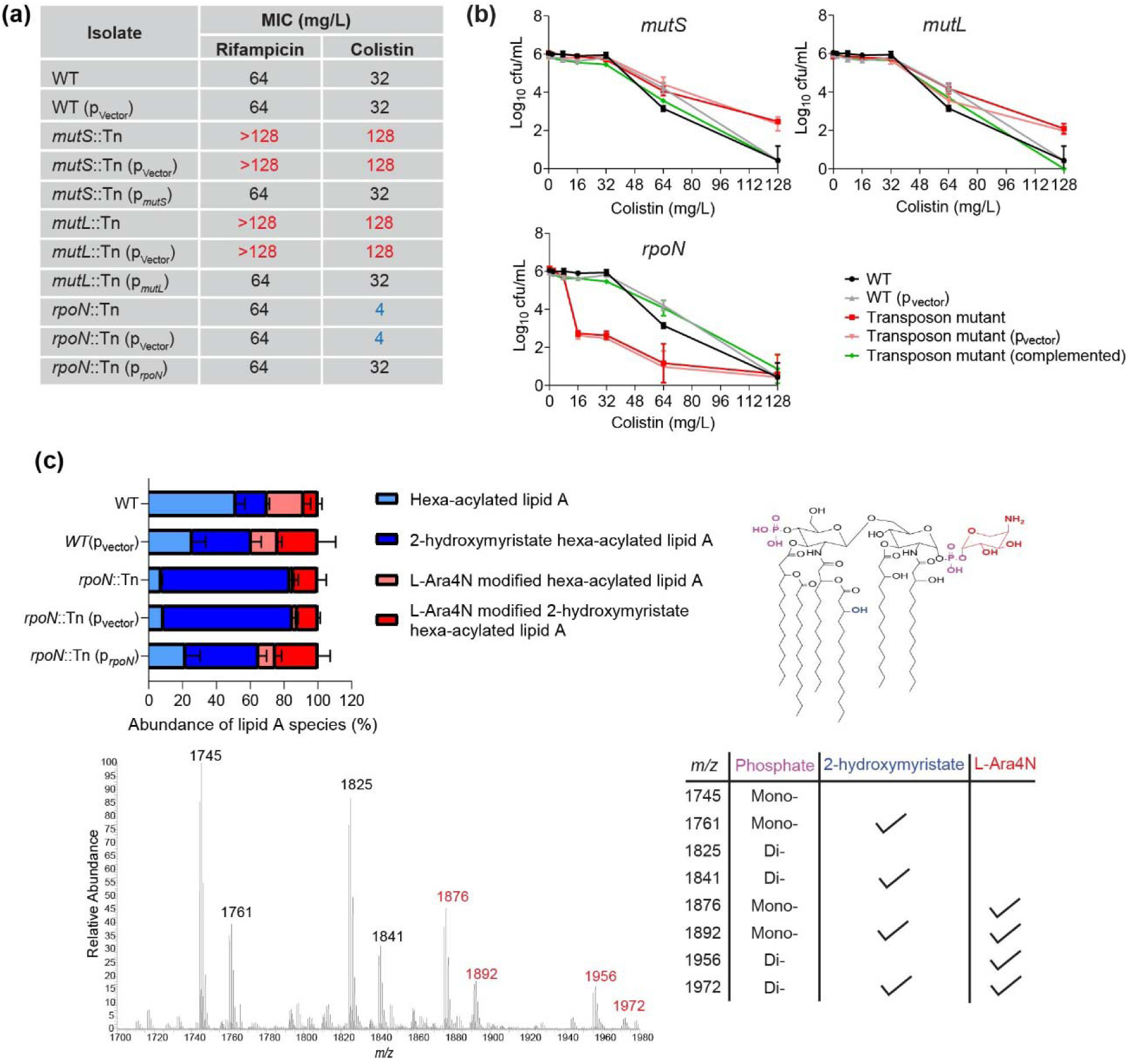
Antibiotic susceptibility and lipid A profiles. **(a)** Minimum inhibitory concentrations (MICs) of rifampicin and colistin. Increased and decreased MICs compared to *K. pneumoniae* MKP103 (WT) were represented by red and blue font colors, respectively. **(b)** Population analysis profiles of *K. pneumoniae* strains on MH agar containing colistin at 2, 4, 8, 16, 32, 64 and 128 mg/L. Data are presented as mean ± SD (*n* = 3). **(c)** Analysis of the individual lipid A species abundance relative to the total abundance. Data are presented as mean ± SD (*n* = 3). Representative mass spectrum profile and chemical structure of each lipid A species.

## Discussion

With the prospect of utilizing phages and antibiotics as effective therapeutic combinations, there have been a number of reports providing evidence on the increased antibiotic susceptibility following phage treatment. These include increased bacterial susceptibility to beta-lactams of a phage-resistant *A. baumannii* due to capsule loss (15), and to erythromycin, tetracycline and ciprofloxacin of *Pseudomonas aeruginosa* treated with phage OMKO1 (known to interact with efflux pumps) (14). In *K. pneumoniae*, the loss of the MDR phenotype has also been reported for the phage-resistant population, caused by the loss of antibiotic resistance gene cassettes [*bla_CTX--M_*, *ant(3”)*, *sul2*, *folA*, *mph(E)/mph(G)*] from a plasmid (21). In the present study, we discovered a bidirectional antibiotic susceptibility effect caused by different phage resistance mechanisms.

The majority phage resistance studies in the literature have employed population-based whole-genome sequencing methods, revealing the predominant phage resistance mechanisms, most often are those associated with phage binding sites such as capsule, lipopolysaccharide (LPS) and outer membrane proteins (15, 23–25). Utilizing genome-wide transposon screening, in addition to the disrupted phage binding site (*fhuA*::Tn and *tonB*::Tn), our study identified phage resistance mechanisms associated with an increased bacterial mutation frequency to phage (*mutS*::Tn and *mutL*::Tn) and a prolonged phage latent period (*rpoN*::Tn and *mnmE*::Tn). Importantly, these mutants demonstrated varying degrees of phage resistance (**Figure 2b and 2c**), suggesting the potential differential mechanisms affecting the phage infectivity.

Hypermutators, characterized by dysfunctional *mutS* and *mutL*, have been frequently associated with antibiotic resistance (26). Among 38 antibiotics examined, *K. pneumoniae* MKP103 with disrupted *mutS* or *mutL* demonstrated increased MICs of colistin and rifampicin. Worth noting, the association of hypermutator phenotype with phage resistance remains relatively scarce. Our findings show that while the plaque formation ability of pKMKP103_1 was unaffected when plated on agar, the lower bacterial reduction observed in the time-kill kinetics for *mutS*::Tn and *mutL*::Tn was likely due to enhanced regrowth dynamics of resistant mutant populations in a liquid culture environment (**Figure 2b and 2c**). This underscores the crucial role of the MutS and MutL DNA mismatch repair system in preventing the rapid emergence of phage resistance during its therapy (**Figure 3d**). MutL was reported to induce phage resistance in *P. aeruginosa* by promoting the deletion of a large chromosomal fragment containing *galU*, leading to a lack of *O*-antigen, the binding site of phage PaoP5 (27). On the contrary, we observed here that MutL played a preventive role against phage resistance in *K. pneumoniae* by reducing the bacterial mutation rate to phage pKMKP103_1. These results highlight the multifaceted functionality of the DNA mismatch repair system in relation to phage resistance across diverse phages and bacterial species. Further investigations are warranted to elucidate the genomic changes of these hypermutators that are directly associated with the change of bacterial susceptibility towards phages. Collectively, *K. pneumoniae* with a hypermutator phenotype must be dealt with great caution due to the potential for the emergence of resistance to both phages and antibiotics, rendering both treatments ineffective.

The *rpoN* encodes an RNA polymerase factor sigma-54 which regulates the transcription of genes across diverse cellular functions (28–31). While *rpoN* was previously reported to be associated with phage resistance in *P. aeruginosa*, the specific mechanism underlying its involvement in phage infection remained elusive (32). Our study discovered that the disruption of *rpoN* in *K. pneumoniae* prolonged the latent period of pKMKP103_1, suggesting the potential transcriptional role of RpoN for efficient phage reproduction and this warrants further investigations.

For the first time, we found that the disruption of *rpoN* in *K. pneumoniae* led to a reduced abundance of L-Ara4N-modified lipid A, thereby restoring the bacterial susceptibility to colistin. Resistance to polymyxins (*i.e.,* colistin and polymyxin B) has been commonly conferred by lipid A modifications with positively-charged moieties such as L-Ara4N, regulated by two-component systems, PhoPQ and PmrAB (22). While RpoN has been suggested to be involved in polymyxin resistance independent of two-component systems in *Salmonella* Typhimurium, the mechanism was unclear (33). Our study sheds light on a crucial research direction to uncover the mechanism played by RpoN in polymyxin resistance, elucidating the role of RpoN in lipid A modifications with L-Ara4N.

Excitingly, our study is the first to document the involvement of bacterial *mnmE* in phage infection, in which this gene encodes a GTPase that plays a key role in the modification of tRNA wobble uridine (34). We demonstrated that the disrupted *mnmE* substantially extended the latent period of pKMKP103_1, thereby leading to resistance. Having an unaffected adsorption profile (**Figure 3b**), we are confident that the reduced infectivity of pKMKP103_1 (**Figure 2b**) was due to the compromised capability to produce phage progeny rather than the inability to adsorb to the bacteria. However, conclusive proof on the direct involvement of bacterial *mnmE* in phage replication requires further in-depth mechanistic investigations.

The most common phage resistance mechanisms involve alterations of bacterial surface structures, thus inhibiting phage adsorption (35). While an earlier study revealed that *K. pneumoniae* MKP103 gained resistance to phages Pharr, LKpNIH-2 and LKpNIH-10 via the loss of capsule, altered LPS or OmpC, and disrupted FhuA, respectively (23), our resistome screening pipeline did not identify transposon mutants of genes associated with biosynthesis of capsule, LPS and OmpC. We have proven TonB-dependent FhuA as an essential receptor binding site for pKMKP103_1. The FhuA gating loop has been well characterized as the binding site for phages T1, T5 and Φ80 to *Escherichia coli*, irrespective of conformational changes induced by TonB (36–38). In contrast to these earlier studies involving TonB-independent phages, our study reveals higher binding specificity of pKMKP103_1 to *K. pneumoniae* MKP103, as the disruption of either *fhuA* or *tonB* alone was sufficient to prevent phage binding.

Overall, this study unveils the multiple phage resistance mechanisms that can be gained by *K. pneumoniae* and demonstrates that these mechanisms can shift the bacterial antibiotic susceptibility bidirectionally (**Figure 5**). Our findings showcase the potential of both synergistic and antagonistic effects when phages and antibiotics are combined as therapeutic strategies. Therefore, we cannot simply assume that phage steering always leads to increased antibiotic susceptibility. Garnering comprehensive mechanistic insights into phage resistance and the resulting steering effect represents the key towards designing effective phage-based therapies, particularly when combined with antibiotics.

**Figure 5:**
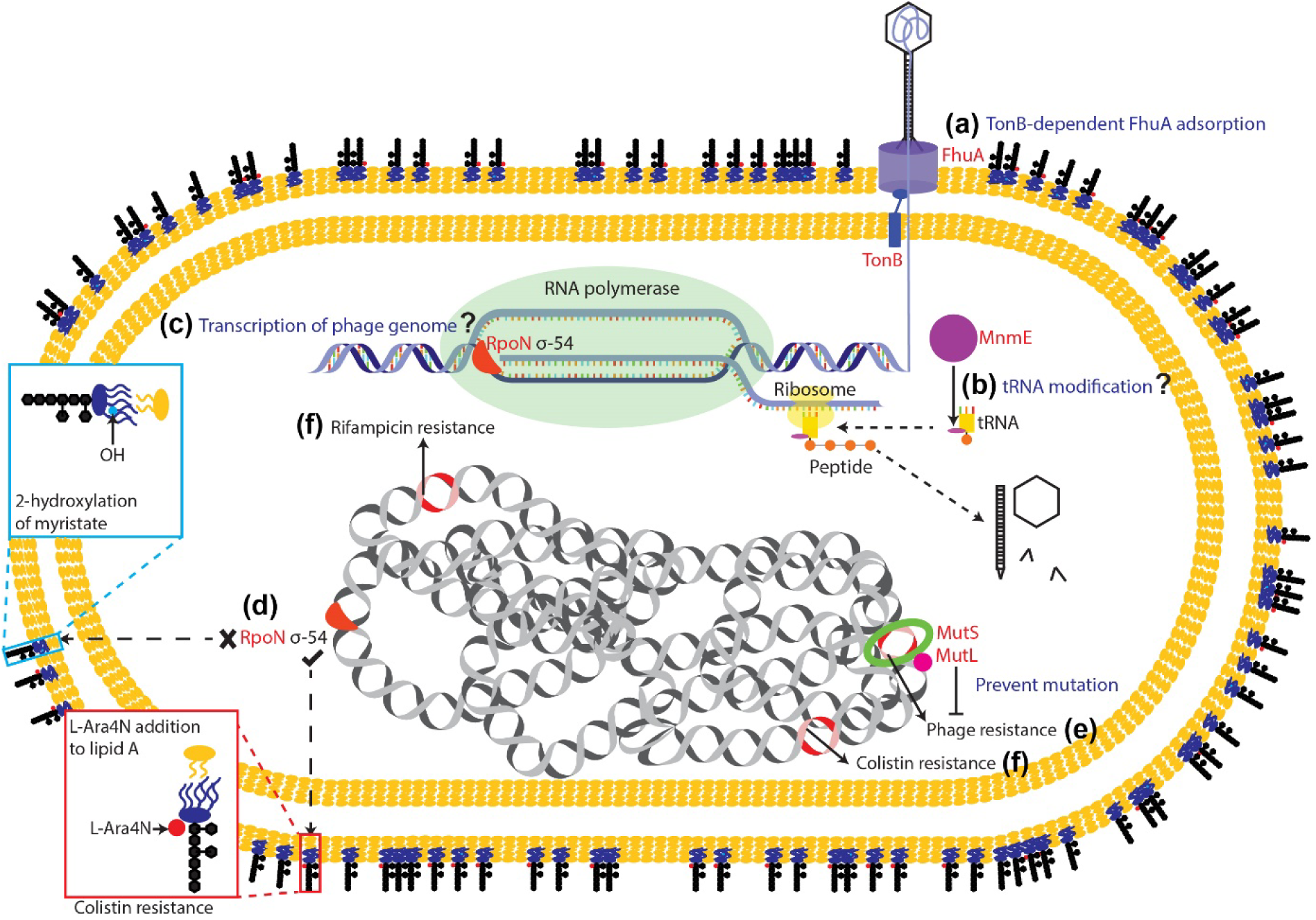
Schematic diagram representing the proposed mechanisms of phage resistance in *K. pneumoniae* MKP103, and the interrelationships with rifampicin and colistin resistance. **(a)** A TonB-dependent FhuA is required for the adsorption of phage. **(b)** MnmE potentially involves in the modification of tRNA, enhancing the protein synthesis for phage particles. **(c)** RpoN could affect the transcription of the phage genome. **(d)** RpoN regulates bacterial transcription for cationic lipid A modifications that confer colistin resistance. MutS and MutL mediate DNA mismatch repair and prevent chromosomal mutations associated with resistance to **(e)** phage, **(f)** rifampicin and colistin.

## Methods

### Bacteria and phage

*K. pneumoniae* MKP103 and its transposon library mutants were used in this study (16). Transposon mutants were selected from Luria Bertani (LB) agar containing 100 mg/L chloramphenicol during initial isolation (16). All experiments were conducted in nutrient broth (Oxoid) unless stated otherwise. Phage pKMKP103_1 was isolated from sewage water obtained from a water treatment plant in Melbourne (Victoria, Australia) with *K. pneumoniae* MKP103 as the host strain. The isolation, purification and amplification of phages were conducted according to the Phage on Tap protocol (39).

### Phage DNA extraction and genome sequencing

Phage pKMKP103_1 was propagated to a titer of ∼10^10^ pfu/mL using the plate lysate method and DNA was extracted using the phenol-chloroform method (39, 40). Briefly, the phage was treated with DNase I (Qiagen) and RNase A (Roche) to remove exogenous bacterial DNA and RNA prior to the addition of sodium dodecyl sulphate and proteinase K to release the DNA from phage particles. Phage DNA was isolated using phenol-chloroform followed by precipitation with isopropanol and 3.0 M sodium acetate (pH 5.2). After two washes with 70% ethanol, DNA was resuspended in DNase-free water. The quality and quantity of DNA were determined using Nanodrop. Sequencing was performed by Genewiz (Suzhou, China) on the Illumina HiSeq. Raw reads (150 bp paired-end) were trimmed and used for *de novo* assembly using SPAdes 3.15.3 (41). Annotation was performed using RAST (42) and PHASTER (43), and hypothetical proteins were further annotated using Blastp (≥90% coverage and ≥80% similarity). Blast Ring Image Generator (BRIG) was used to create a representative visualization of the phage genome (44).

### Transmission electron microscopy

High purity pKMKP103_1 was prepared using the cesium chloride gradient method (45). For TEM imaging, 5 μL of the phage solution was applied to a glow-discharged carbon-coated copper grid and stained with 1% uranyl acetate. Imaging was conducted on a JEOL JEM-1400Plus TEM (Ramaciotti Centre for Cryo-Electron Microscopy, Monash University). ImageJ was used for visualizing and measuring the dimensions of the phage (46).

### Genome-wide transposon screening of phage-resistant mutants

The transposon mutant library of *K. pneumoniae* MKP103 was utilized for the screening of phage pKMKP103_1-resistant mutants (16). A 96-pin replicator was used to inoculate individual mutants into 96-well plates (200 μL of nutrient broth per well) which were then incubated at 37°C for 18 h to reach ∼10^9^ cfu/mL. A 10-fold dilution (20 μL in 180 μL) was conducted twice using 96-well plates to achieve a starting bacterial inoculum of ∼10^7^ cfu/mL. Phage pKMKP103_1 was added to the wells to achieve a multiplicity of infection (MOI) of 1 (∼10^7^ pfu/mL). *K. pneumoniae* MKP103 (WT) was included as the control in each round of the screening experiments. OD_600nm_ was measured at 8 and 10 h with a SPECTROstar^®^ Nano Microplate Reader (BMG Labtech); these time-points were chosen based on preliminary experiments conducted with the WT which demonstrated an increase in OD_600nm_ after 10 h of incubation. Consequently, mutants with greater OD_600nm_ in comparison to the WT at 8 and/or 10 h (OD_600nm_ ≥0.12) were potential resistant mutants. This transposon mutant library consists of 1 to 6 individual transposon-inserted mutant(s) of each non-essential gene (16). The screening process was narrowed to the genes of which ≥2 individual transposon mutants were identified. Additionally, sole transposon mutants were also included to prevent missing out of any potential candidates. With these selected mutants, turbidimetric assays were conducted in three biological replicates followed by time-kill analyses to confirm the phage-resistant phenotype.

### Complementation of mutants

Sanger sequencing was first conducted to confirm that the candidate mutants contain the transposon at the indicated site (**Table S3**). To complement the gene into the respective transposon mutant, the gene was amplified by PCR from the WT (**Table S3**). PCR products and pBBR1MCS-5 carrying the gentamicin resistance cassette (47) were digested over 4 h with the restriction enzymes targeting the relevant restriction sites (**Table S3**). Following clean-up of the digested products, the PCR product and pBBR1MCS-5 were ligated overnight at 4°C. The ligation mixture was then transformed into chemical-competent *E. coli* DH5α and plated on LB agar containing 20 mg/L gentamicin. Colony PCR was conducted to identify the colonies with successful ligation of the gene of interest onto pBBR1MCS-5. The plasmid was extracted from *E. coli* DH5α using a QIAprep^®^ Miniprep Kit and Sanger sequencing was conducted to confirm the correct cloning. The complemented plasmids were electroporated into the electrocompetent transposon mutants of the respective gene. An empty pBBR1MCS-5 vector (p_Vector_) was also electroporated into the WT and all mutants as control strains to exclude vector effects. Successful transformation was confirmed with the growth of colonies on LB agar containing 20 mg/L gentamicin, followed by PCR.

### Turbidimetric assay

*K. pneumoniae* strains were grown in nutrient broth at 37°C with shaking (200 rpm) for 18 h. The overnight bacterial culture was inoculated in fresh nutrient broth (1:100 dilution) and incubated until a log-phase with an OD_600nm_ of 0.5 (equivalent to ∼10^8^ cfu/mL) was achieved. The culture was then diluted 1:10 with fresh nutrient broth to reach a final inoculum of ∼10^7^ cfu/mL. Phage pKMKP103_1 was added at an MOI of 1 (final titer of ∼10^7^ pfu/mL) or 10 (final titer of ∼10^8^ pfu/mL). Continuous OD_600nm_ readings were taken hourly up to 20–24 h with a SPECTROstar^®^ Nano Microplate Reader (BMG Labtech).

### Time-kill kinetics

*K. pneumoniae* strains were grown in nutrient broth at 37°C with shaking (200 rpm) for 18 h. The overnight bacterial culture was inoculated in fresh nutrient broth (1:100 dilution) and incubated until a log-phase with an OD_600nm_ of 0.5 (equivalent to ∼10^8^ cfu/mL) was achieved. The culture was then diluted 1:10 with fresh nutrient broth to reach a final inoculum of ∼10^7^ cfu/mL. Phage pKMKP103_1 was added at an MOI of 10 (final titer of ∼10^8^ pfu/mL). For mutants carrying the empty vector and complemented strains, 5 mg/L gentamicin was added as the selection pressure to maintain the plasmid throughout the experiment. An aliquot of 200 μL of culture was collected at 0, 1, 2, 4 and 24 h, centrifuged at 10,000 × *g*, and the bacterial pellet was resuspended in an equal volume of 0.9% sodium chloride (NaCl). Following 10-fold dilutions with 0.9% NaCl, 10-μL spots were plated on nutrient agar for viable counting.

### Efficiency of plating (EOP)

The EOP was conducted with modifications (48). An overnight bacterial culture was prepared as described above, with 100 μL of the culture subsequently mixed with 4 mL soft agar and poured onto a nutrient agar plate to form double-layered agar containing bacteria. Phage pKMKP103_1 at 10^9^ pfu/mL (determined with the WT) was subjected to a series of 10-fold dilutions using SM buffer (50 mM Tris-HCl, 100 mM NaCl, 8 mM MgSO_4_, pH7.4) and plated onto the double-layered agar as 10-μL spots. Plates were then incubated overnight at 37°C before enumeration of visible plaque.

### Phage adsorption assay

*K. pneumoniae* was grown to log-phase as described above and diluted to ∼10^7^ cfu/mL (final volume of 1 mL). Phage pKMKP103_1 was then added to reach a final titer of 2.5 × 10^4^ pfu/mL (MOI of 0.0025). At 2, 4 and 10-min following co-incubation, 100 μL of culture was collected and centrifuged at 16,000 × *g* for 1 min. The supernatant containing the unabsorbed phage was then spotted in three technical replicates on double-layered agar containing either *K. pneumoniae* MKP103 or *K. pneumoniae* MKP103 (p_Vector_); strains carrying the plasmid (empty vector and complemented) were plated with *K. pneumoniae* MKP103 (p_Vector_). Plaques were enumerated following overnight incubation at 37°C. Three biological replicates were conducted for each group.

### Fluorescent and confocal microscopic assay

Phage pKMKP103_1 was incubated with 100× SYBR™ Gold Nucleic Acid stain (Invitrogen™) at 4°C for 4 h. The stained pKMKP103_1 was then transferred to an Amicon^®^ Ultra-15 centrifugal filter tube (molecular weight cut-off 100 kDa) and centrifuged at 3,220 × *g* (4°C) to remove excess stain. Two further washes with SM buffer were conducted prior to resuspension of the stained pKMKP103_1 in an equal volume of SM buffer. Enumeration of the plaques was performed to determine the titer of stained pKMKP103_1 for the microscopic assays. The log-phase bacterial culture was grown and diluted to 2 × 10^7^ cfu/mL, and incubated with FM™ 4-64FX membrane stain (Invitrogen™) at room temperature for 5 min. The SYBR™ Gold stained pKMKP103_1 was added to the FM™ 4-64FX stained *K. pneumoniae* at an MOI of 100 (fluorescent microscopy) and 1,000 (confocal microscopy). For fluorescent microscopy, the mixture was applied directly onto the glass slide. For confocal microscopy, the mixture was mixed with molten 2% agarose (55°C) to fix the sample for imaging.

### One-step growth curve

Log-phase *K. pneumoniae* culture was diluted to ∼10^7^ cfu/mL and pKMKP103_1 was added to achieve a final titer of 10^5^ pfu/mL (MOI 0.01). Following 2-min incubation at 37°C the culture was centrifuged at 16,000 × *g* for 1 min, the supernatant containing the free phage was removed and the bacterial pellet was resuspended in an equal volume of nutrient broth. Aliquots (100 μL) were then removed at 10-min intervals and added to a glass tube containing 4 mL soft agar with 100 μL bacterial culture of *K. pneumoniae* MKP103 or *K. pneumoniae* MKP103 (p_Vector_).

Following a rapid vortex, the entire content of the tube was poured onto a nutrient agar plate to form a double-layered agar. Plaques were enumerated following overnight incubation at 37°C.

### Mutation frequency to pKMKP103_1

The mutation frequency of *K. pneumoniae* strains to pKMKP103_1 was examined using the agar overlay method with modifications (49). *K. pneumoniae* was grown to log-phase as described above to an OD_600nm_ of 0.5 (equivalent to ∼10^8^ cfu/mL). For mutants carrying the empty vector and complemented strains, 5 mg/L gentamicin was added as the selection pressure to maintain the plasmid. Bacterial culture was centrifuged at 16,000 × *g* at 4°C and the pellet was resuspended in an equal volume of 0.9% NaCl. The bacterial culture was serially diluted and 100 μL was added to a glass tube containing 4 mL soft agar with pKMKP103_1 (10^9^ pfu/mL). Control samples without phage were included to obtain the total number of colonies. A double-layered agar was prepared as described above. Colonies were counted following a 20-h incubation at 37°C. Mutation frequency was calculated by dividing the number of colonies formed on the agar containing pKMKP103_1 over the total number of colonies. Statistical analysis was conducted using the multiple comparisons with uncorrected Dunn’s test on GraphPad Prism 9.3.1.

### Antibiotic susceptibility testing and population analysis profiles (PAPs)

Susceptibility of *K. pneumoniae* strains to a range of antibiotics was determined using broth microdilution (Trek Sensititre^®^, ThermoFisher Scientific) as per manufacturer’s instructions. The lowest antibiotic concentration that inhibited the visible growth of bacteria was the MIC. For mutants that showed a 2-fold difference in MIC compared to the WT and demonstrated restoration of the MIC following complementation (**Table S2**), additional broth microdilution assays were conducted in two technical replicates and confirmed on two separate occasions. All MIC assays were conducted in cation-adjusted (22.5 mg/L Ca^2+^, 11.25 mg/L Mg^2+^) Mueller Hinton broth (CAMHB). PAPs were undertaken as described previously with minor modifications (50). Briefly, bacterial culture was grown in CAMHB until log-phase and following 10-fold serial dilutions, cultures were plated onto MH agar containing colistin at 2, 4, 8, 16, 32, 64 or 128 mg/L. Viable colonies were enumerated following overnight incubation at 37°C.

### Lipid A profiling

A single bacterial colony was inoculated in CAMHB and incubated at 37°C with shaking (200 rpm) for 18 h. The overnight culture was then diluted 1:100 with fresh CAMHB (final volume, 200 mL) and grown to an OD_600nm_ of ∼0.7 (8.35 ± 0.13 log_10_ cfu/mL). Bacterial pellets were collected via centrifugation at 9,000 × *g* for 10 min, washed twice with 0.9% NaCl, and resuspended in 4 mL of 0.9% NaCl. Lipid A extraction was conducted with minor modifications (51). Methanol (10 mL) and chloroform (5 mL) were then added to prepare a single-phase Bligh-Dyer mixture (chloroform:methanol:water, 1:2:0.8, v/v) for the extraction of LPS. LPS pellet was obtained by centrifugation at 3,220 × *g* for 10 min and washed once with single-phase Bligh-Dyer mixture. The washed pellet was resuspended in 5.4 mL of hydrolysis buffer (50 mM sodium acetate, pH 4.5) and incubated in a boiling water bath for 1 h. Following hydrolysis, methanol (6 mL) and chloroform (6 mL) were added to make a two-phase Bligh-Dyer mixture (chloroform:methanol:water, 1:1:0.9, v/v). Following centrifugation at 3,220 × *g* for 10 min, the lower phase of the mixture containing the lipid A was collected, dried and stored at -20°C. Prior to LC-MS analysis, the dried lipid A was resuspended in 200 μL of chloroform:methanol (1:1, v/v), centrifuged at 14,000 × *g* for 10 min, and 150 μL of supernatant was collected. An equal volume of isopropanol:water (2:1, v/v) was then added prior to centrifugation at 14,000 × *g* for 10 min, and 200 μL of supernatant was obtained for LC-MS analysis on the Dionex U3000 high-performance liquid chromatography (HPLC) system in tandem with a Q-Exactive Orbitrap high-resolution mass spectrometer (Thermo Fisher).

## Author contributions

Conceptualization, S.C.N. and J.Li; Methodology, S.C.N., M.A.K.A., M.H., Y.Z. and D.T.M.; Investigation, S.C.N., J.Lu, H.H.Y., H.W., M.A.K.A., M.H., J.Z., N.S., B.P.H., Y.Z. and D.T.M.; Writing – Original Draft, S.C.N.; Writing – Review & Editing, S.C.N., P.J.B., T.V. and J.Li; Funding acquisition, G.R., T.V., S.A., R.T.S., J.J.B. and J.Li; Supervision, G.R., T.V., S.A., R.T.S., J.J.B. and J.Li.

## Acknowledgements

This study was funded by the National Institute of Allergy and Infectious Diseases of the National Institutes of Health grant (R21 AI156766) and Monash-UCSD Seed Fund. J.Li is a National Health and Medical Research Council (NHMRC) Principal Research Fellow (APP1157909). The content is solely the responsibility of the authors and does not necessarily represent the official views of the National Institute of Allergy and Infectious Diseases or the National Institutes of Health.

We would like to express our gratitude to the Ramaciotti Centre for Cryo-Electron Microscopy (Monash University) and Micromon Genomics (Monash University) for their facilities to conduct TEM imaging and Sanger sequencing.

