## Supplementary material for "Diversified repertoire of phage resistance in *Klebsiella pneumoniae* and bidirectional steering effects impacting antibiotic susceptibility": Table S3

**Table S3:** Primers used in this study.

| Confirmation of transposon mutants | | |
|---|---|---|
| **Name** | **Sequence (5' – 3')** | **Purpose** |
| *mutS*_555_Fw | CGAATCGTCGCTGATCGAGG | Confirmation of *mutS* transposon mutant |
| *mutS*_2022_Rv | CATCTCCACCATAAAGGTCGAGC | |
| *mutL*_156_Fw | GCTCATTCGCATCCGCGATAAC | Confirmation of *mutL* transposon mutant |
| *mutL*_1140_Rv | CGATGGCGTGATGCTGAAGC | |
| *fhuA*_190_Fw | GCAAAACGCTCCGCCACGG | Confirmation of *fhuA* transposon mutant |
| *fhuA*_2034_Rv | CAGCGCATAACCGGCCACAC | |
| *tonB*_4_Fw | GTTAATCTCGACGCCGTTCAGG | Confirmation of *tonB* transposon mutant |
| *tonB*_+5_Rv | CGACTATGAGCGCAATGACCC | |
| *rpoN*_250_Fw | CTCGATGCCAGCTGGGATGA | Confirmation of *rpoN* transposon mutant |
| *rpoN*_1324_Rv | GCTTACTGTCGCTCAACGGC | |
| *mnmE*_3_Fw | GAGCCATAACGACACTATCGTCG | Confirmation of *mnmE* transposon mutant |
| *mnmE*_1140_Rv | GGTATCGAAGCCCATGCTCTG | |
| Pcm-140 | CTGCGAAGTGATCTTCCGTCAC | Transposon-specific primer |

| Complementation of mutants | | | |
|---|---|---|---|
| **Name** | **Restriction site** | **Sequence (5' – 3')** | **Purpose** |
| *mutS*_BamHI_F | BamHI | CACCGTggatccCATGAGCACAATTGACAATC | Cloning of *mutS* |
| *mutS*_XbaI_R | XbaI | GGTAGCtctagaTTCACCTTAATGGGGGGCT | |
| *mutL*_BamHI_F | BamHI | CACCGTggatccGATGCCGATTCAGGTTCT | Cloning of *mutL* |
| *mutL*_XbaI_R | XbaI | GGTAGCtctagaCATCATTCATCTTTCAGGG | |
| *fhuA*_HindIII_F | HindIII | CACCGTaagcttCATGGCGCGTCCAAAAAC | Cloning of *fhuA* |
| *fhuA*_XbaI_R | XbaI | GGTAGCtctagaGGTTAGTAACGGAAGGTG | |
| *tonB*_BamHI_F | BamHI | CACCGTggatccCTATGAGCGCAATGAC | Cloning of *tonB* |
| *tonB*_XbaI_R | XbaI | GGTAGCtctagaATCAGTTAATCTCGACGC | |
| *rpoN*_HindIII_F | HindIII | CACCGTaagcttCATGAAGCAAGGTTTGC | Cloning of *rpoN* |
| *rpoN*_XbaI_R | XbaI | GGTAGCtctagaCGGTTATGTCAGACCAG | |
| *mnmE*_BamHI_F | BamHI | CACCGTggatccCATGAGCCATAACGACA | Cloning of *mnmE* |
| *mnmE*_XbaI_R | XbaI | GGTAGCtctagaGTTATTTGCCGATACAGAAG | |

| | | | |
|---|---|---|---|
| pBBR1MCS_Fw | - | AACGACGGCCAGTGAGCG | Confirmation of cloning |
| pBBR1MCS_Rv | - | GAACAAAAGCTGGGTACCG | |
